# Genetic Foundations of Inter-individual Neurophysiological Variability

**DOI:** 10.1101/2024.07.19.604292

**Authors:** Jason da Silva Castanheira, Jonathan Poli, Justine Y. Hansen, Bratislav Misic, Sylvain Baillet

**Affiliations:** Montreal Neurological Institute, McGill University, Montreal QC, Canada; CentraleSupélec, Université Paris-Saclay, Paris, France

**Keywords:** Neurophysiology, Heritability, Genetic determinants, Brain-fingerprints, Magnetoencephalography (MEG), Monozygotic and dizygotic twins, Neurotransmission, Neurodevelopment, Cognitive and affective brain functions

## Abstract

Neurophysiological brain activity shapes cognitive functions and individual traits. Here, we investigated the extent to which individual neurophysiological properties are genetically determined and how these adult traits align with cortical gene expression patterns across development. Using task-free magnetoencephalography in monozygotic and dizygotic twins, as well as unrelated individuals, we found that neurophysiological traits were significantly more similar between monozygotic twins, indicating a genetic influence, although individual-specific variability remained predominant. These heritable brain dynamics were predominantly associated with genes involved in neurotransmission, expressed along a topographical gradient that mirrors psychological functions, including attention, planning, and emotional processes. Furthermore, the cortical expression patterns of genes associated with individual differentiation aligned most strongly with gene expression profiles observed during adulthood in previously published longitudinal datasets. These findings underscore a persistent genetic influence on neurophysiological activity, supporting individual cognitive and behavioral variability.

## Main Text

Several recent neuroimaging studies have shown that ongoing brain activity at rest, without performing a specific task, defines a neurophysiological profile unique to each person. Unlike hand fingerprints, these *brain-fingerprints* are associated with individual cognitive traits and are altered by pathology (*1–10*), forming a distinct personal neurophysiological profile. Whether a person’s genotype is associated with their neurophysiological profile is currently unknown. Heritability studies of inter-individual variability (*11*) have reported some genetic associations with brain structures (*12–15*) and activity (*16–21*). Genetic factors determine, to some extent, inter-individual variations in broad cognitive domains such as attentional and general intellectual abilities (*22–26*). Here, we studied how both individual neurophysiological and cognitive profiles relate to the spatial organization of gene expression patterns.

We used task-free magnetoencephalographic (MEG) imaging (*27*) to derive the individual neurophysiological profiles (*2*) of monozygotic (MZ) and dizygotic (DZ) twins, along with unrelated individuals. We hypothesized that if neurophysiological brain activity is determined by genetic factors, then the neurophysiological profiles of MZ twins, who share nearly identical genomes, would be nearly identical, unlike those of DZ twins (*28*). We then identified which genes are related to the features that differentiate individuals based on their neurophysiological profiles. Additionally, we investigated whether the genetic influence on these neurophysiological traits is linked to major psychological processes that characterize individual traits and how this influence aligns with cortical gene expression patterns throughout development.

## Results

### Individual Neurophysiological Profiling and Differentiation

We derived the neurophysiological profile of 89 individuals (17 pairs of monozygotic twins, 11 pairs of dizygotic twins, and 33 unrelated individuals; 22-35 years old) from three 6-minute task-free MEG recordings provided by the Human Connectome Project(*29*). We derived individual profiles from the distribution of neurophysiological signal power of MEG activity across the cortex for each recording (*2*) (see Methods).

We first assessed the accuracy of inter-individual differentiation based on the neurophysiological profiles obtained from the three recordings of each participant (Figure 1). The accuracy of inter-individual differentiation from these neurophysiological profiles was 83.4% (95% bootstrapped CI [73.8, 90.0]; Figure S1) across a broad frequency spectrum of brain activity (1-150 Hz). The high temporal resolution of the data enabled us to study how the accuracy of neurophysiological profiling varied between the prototypical frequency bands of electrophysiology. Inter-individual differentiation varied substantially across these frequency ranges, from 59.7% in the delta band (1-4 Hz; [57.5, 73.8]) to 87.4% in the high-gamma band (50-150 Hz; [80.0, 92.5]) as detailed in Figure S1.

**Fig. 1.**
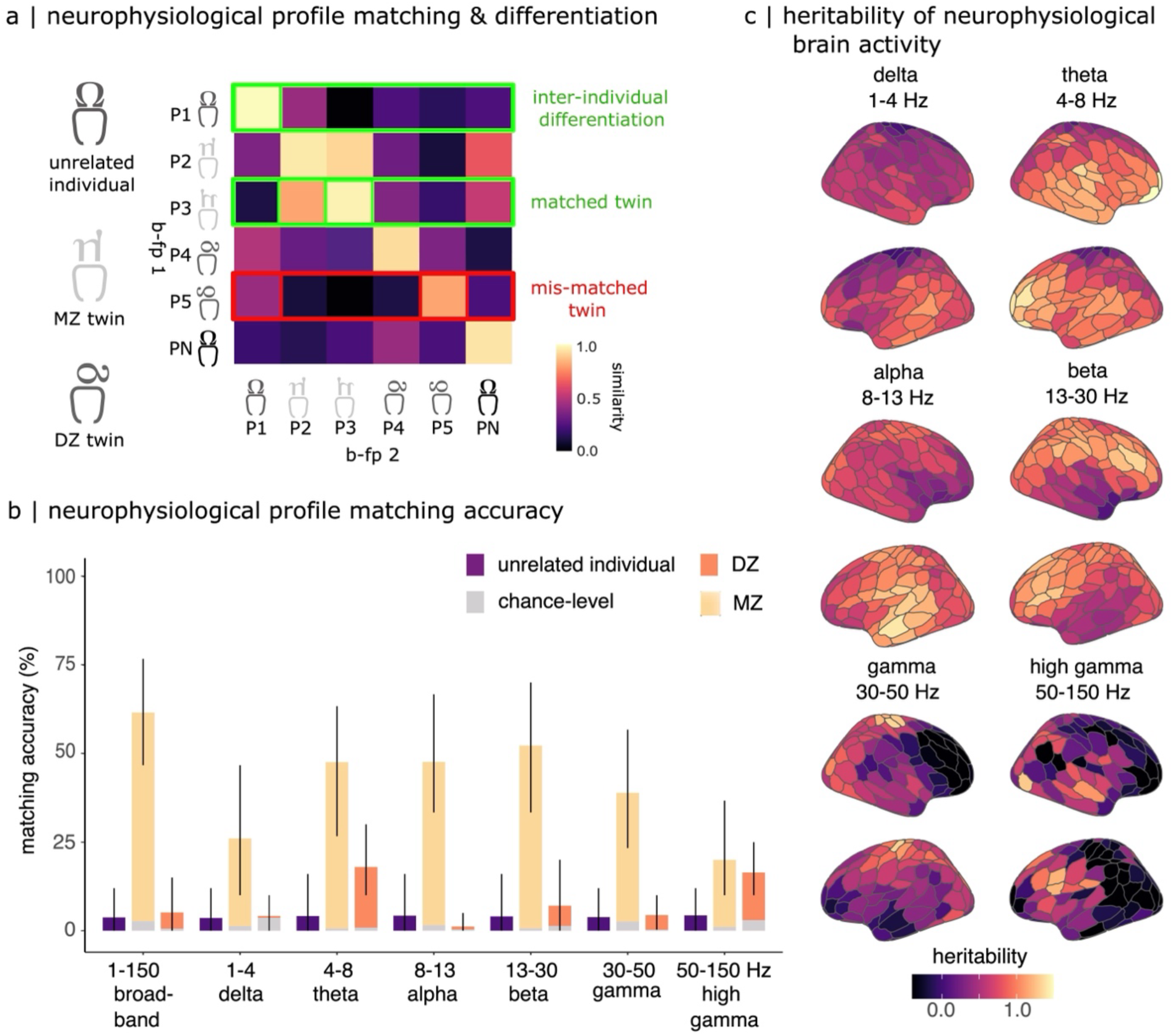
Neurophysiological Profiling. a) Color-coded similarity matrix showing self-similarity (diagonal elements) and between-participant similarity (off-diagonal elements) of neurophysiological profiles, computed across three independent recordings. Similarity was assessed using cross-correlation coefficients. A participant was considered correctly differentiated if their own profiles were more similar across sessions than to profiles from others. Twin sibling matching was assessed using the same similarity statistics. b) Bar plots showing twin pair matching accuracies across broadband (1–150 Hz) and standard electrophysiological frequency bands (delta (1-4 Hz), theta (4-8 Hz), alpha (8-13 Hz), beta (13-30 Hz), gamma (30-50 Hz), and high gamma (50-150 Hz)). Orange bars represent monozygotic (MZ) and dizygotic (DZ) twin matching accuracies; purple bars represent matching accuracies between randomly paired unrelated individuals. Gray bars represent chance-level matching based on mock neurophysiological profiles derived from empty-room MEG recordings. Error bars denote 95% confidence intervals. c) Cortical maps of heritability estimates for neurophysiological traits, computed using Falconer’s equation, illustrating the spatial distribution of genetic contributions across the cortex. Legend: b-fp1: first brain-fingerprint; b-fp2: second brain-fingerprint; MZ: monozygotic twins; DZ: dizygotic twins.

We then evaluated the similarity between the neurophysiological profiles of siblings in a twin pair (Figure 1A), finding that those of monozygotic twins matched with 61.5% accuracy ([46.7, 76.7]; Figure 1B). In contrast, the profiles of dizygotic twins showed a considerably lower match accuracy of only 5.2% ([0.0, 15.0]; Figure 1B). These differences in matching accuracies between monozygotic and dizygotic twins varied across the frequency bands of neurophysiological brain activity, as shown in Figure 1B. The discrepancies were particularly pronounced in the alpha band (8-13 Hz), where MZ twins matched with an accuracy of 47.8% ([33.3, 66.7]) compared to only 1.2% for DZ twins ([0.0, 5.0]). In the beta band (13-30 Hz), accuracies were 52.1% for MZ twins ([33.3, 70.0]) versus 7.1% for DZ twins ([0.0, 20.0]; Figure 1B).

To benchmark chance-level matching performance, we randomly created pairs of unrelated individuals and computed their neurophysiological profile matching accuracies, repeating this procedure 300 times per frequency band. Across all bands, matching unrelated individuals yielded low accuracy (<5%, Figure 1B), comparable to the matching accuracy observed for dizygotic (DZ) twin pairs. In contrast, monozygotic (MZ) twin pairs exhibited significantly higher matching accuracies for broadband (1–150 Hz), theta (4–8 Hz), alpha (8–13 Hz), beta (13–30 Hz), and gamma (30–50 Hz) neurophysiological profiles (Figure 1B). These results confirm that while neurophysiological profiles are more similar within MZ twin pairs, DZ twin similarity is not distinguishable from unrelated individuals.

To ensure the robustness of our results, we computed chance-level neurophysiological profiling and twin pair matching based on environmental and equipment noise using empty-room recordings taken before each MEG recording session (Figure 1B). We observed above-chance differentiation across all frequency bands, including high frequencies, confirming that differentiation effects were not driven by SNR biases. We removed the variance associated with physiological artifacts from the neurophysiological profiles and found that our results remained robust (Figure S2).

Refer to Supplemental Figure S3 for the distributions of self-, other-, and twin-pair similarity between neurophysiological profiles. While differentiation accuracy remains our primary metric, we provide additional analyses illustrating that MZ twins exhibit higher neurophysiological profile similarity than DZ twins, while unrelated individuals do not significantly differ from either group. These findings reinforce that differentiation accuracy is a robust measure of individual distinctiveness in neurophysiology.

### Heritability of Neurophysiological Traits

Although we reported above that the neurophysiological profiles of MZ twins match each other more closely than those of the general population (Figure 1B, yellow bars), the neurophysiological profile of each MZ twin remains distinguishable from their sibling such that we can still correctly differentiate individual twins from a cohort that includes their sibling (Figure S1, purple bars). One possible explanation is that individual-specific features stand apart from the heritable aspects of their neurophysiological profile. We therefore sought to confirm that the most differentiable features of neurophysiological profiles are similarly heritable.

To assess this, we measured the spatial alignment between the most salient features for participant differentiation (Figure S4A-B) and the heritability of neurophysiological traits (Figure 1C & S4C). While absolute heritability values provide an upper bound on genetic influence, our primary focus was on spatial patterns of heritability across frequency bands and cortical regions.

First, we leveraged intra-class correlation (ICC) statistics (*2*, *3*, *30*) to quantify which cortical parcel and frequency band contributed the most toward participant differentiation (see Methods). ICC measures the ratio of within-participant to between-participant variance, where higher ICC values indicate that a neurophysiological trait can robustly differentiate individuals across multiple recordings. Our analysis revealed that posterior cortical activity in the theta (4-8 Hz; ICC = 0.74), alpha (8-13 Hz; ICC = 0.83), and beta (13-30 Hz; ICC = 0.74) frequency bands were the most distinctive traits for individual differentiation (Figure S4B).

Next, we quantified the heritability of these neurophysiological traits using Falconer’s method. We observed mean heritability (H^2^) values of 0.85 for theta-band traits, 0.76 for alpha-band traits, and 0.77 for beta-band traits across the temporal, frontal, and parietal-occipital cortices (Figure 1C). The occipital visual regions showed the highest heritability, whereas the limbic network showed the lowest (Figure 1C).

Permutation tests confirmed significant cortical-wide heritability in the alpha band, as well as regionally significant heritability in frontal and parietal areas for beta-band traits and frontal regions for theta-band traits (Figure S5).

Finally, we observed significant spatial correlations between ICC-derived differentiation maps and heritability maps, confirming that the most distinctive features of neurophysiological profiles tend to be highly heritable. These correlations were significant across broadband cortical signals (1-150 Hz; r = 0.28, pspin = 0.026), with particularly strong relationships in the alpha (r = 0.62, p_spin_ = 0.0009) and beta (r = 0.58, p_spin_ = 0.0009) bands (Figure S6 left panel). These findings confirm that the most distinctive features of neurophysiological profiles tend to be heritable, further emphasizing the genetic basis for the neurophysiological characteristics that distinguish individuals.

### Person-Specific Neurophysiological Signals are Aligned with Cortical Gene Expression

We then assessed whether neurophysiological profiles for participant differentiation (full ICC maps from Figure S4) also align topographically with genetic cortical expressions. To do this, we studied the spatial covariation of the ICC of neurophysiological profiles with maps of cortical gene expressions with greater differential stability (>0.1) (*31–35*), retrieved from the microarray Allen Human Brain Atlas (*33*), using Partial Least Squares (PLS) correlation (Figure 2A). We found a single PLS component significantly accounting for 85.2% of the covariance (CI [73.4, 90.1], p_spin_ = 0.01; Figure 2B). This analysis revealed a topographical pattern where visual and somatomotor regions exhibited positive covariance with genetic expressions, while limbic regions exhibited negative covariance (Figure 2B). This indicates that the most differentiable traits of individual neurophysiological profiles are spatially aligned with specific gene expression patterns along the cortical surface. We cross-validated this observation with 1,000 permutations corrected for spatial autocorrelation, resulting in a median out-of-sample correlation of r = 0.64 (p_spin_ = 0.002; Figure 2C).

**Figure 2:**
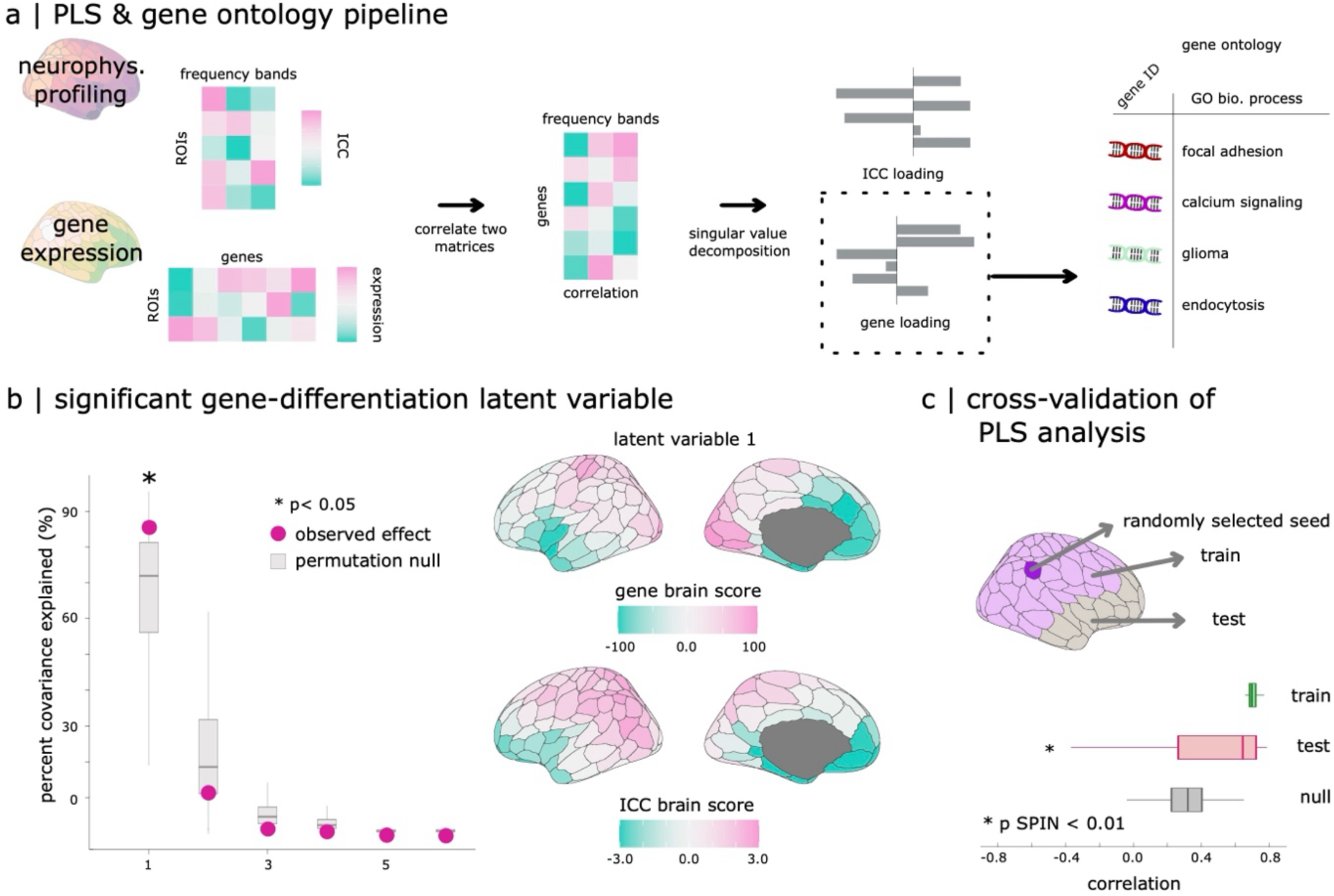
Analysis Pipeline and Outcomes of Gene-Differentiation PLS Analysis. a) Two data matrices were submitted to a Partial-Least-Squares (PLS) and gene ontology analysis: i) the first data matrix gathered the most salient traits for neurophysiological profiling; ii) the second data matrix contained scores of gene expression across the regions of the Schaefer-200 cortical atlas(*36*). The PLS analysis resulted in latent components capturing the modes of largest covariance between these variables. Using the elements with top loadings, we performed a gene ontology analysis to determine if the contributing genes were enriched for specific molecular processes. b) The left panel shows the PLS latent components with pink dots, ordered by decreasing effect size. Statistical significance was determined with 1,000 permutations of the observed data, with spatial autocorrelation correction applied, highlighting only the first latent component. The right panel shows the related cortical topographies of gene-expression and ICC scores derived by projecting this first latent component onto the observed data. Positive gene brain scores positively covary with ICC brain scores, and negatively covary with negative ICC brain scores. c) We trained the PLS model using 75% of the cortical regions, selected based on their proximity in Euclidean distance to a randomly selected seed (dark purple regions), and tested the relationship between gene-expression and ICC scores on the rest of the data. The median out-of-sample relationship observed was r= 0.64 (p_spin_= 0.002).

We then measured how each frequency range of cortical activity contributed to this alignment between cortical gene expression and the salient traits of neurophysiological profiles. To do so, we computed the loadings for each frequency band, which correspond to the Pearson correlation coefficients between ICC data and the observed cortical score pattern (see Methods). We found that all frequency ranges contributed exclusively positively to this association: delta (1-4 Hz; r = 0.52, 95% CI [0.45, 0.61]), alpha (8-13 Hz; r = 0.63, 95% CI [0.58, 0.69]), beta (13-30 Hz; r = 0.52, 95% CI [0.42, 0.62]), gamma (30-50 Hz; r = 0.71, 95% CI [0.66, 0.76]), and high gamma bands (50-150 Hz; r = 0.43, 95% CI [0.34, 0.53]) with the exception of the theta band (4-8 Hz; r = 0.07, 95% CI [-0.09, 0.22]).

### Genes and Cell Types Associated with Neurophysiological Traits

We then investigated which genes’ expressions contribute the most to the reported association with neurophysiological traits, aiming to identify the biological functions associated with these genes and the specific cell types involved. We selected the top 2,208 genes based on their highest positive loadings in the PLS analysis and the top 2,344 genes based on their highest negative loadings. We performed a gene ontology (GO) analysis using the ShinyGO pipeline and resources from the GO database (*37*), which revealed distinct biological processes linked to these sets of genes (see Methods).

Genes with positive loadings were enriched in biological processes such as ion transport, synaptic functioning, and neurotransmitter release, while genes with negative loadings were associated with processes like development, neurogenesis, and cell morphogenesis (Figure 3A, bottom panel; complete gene list in Supplemental Information). In short, positively weighted genes covary positively with participant differentiation (i.e., the positive ICC brain scores) which in turn is associated with neurochemical signalling. Additionally, we verified that the spatial alignment between gene-ontological categories was not driven by the spatial autocorrelation of cortical maps through SPIN tests (colours of points in Figure 3A; see Methods).

**Figure 3:**
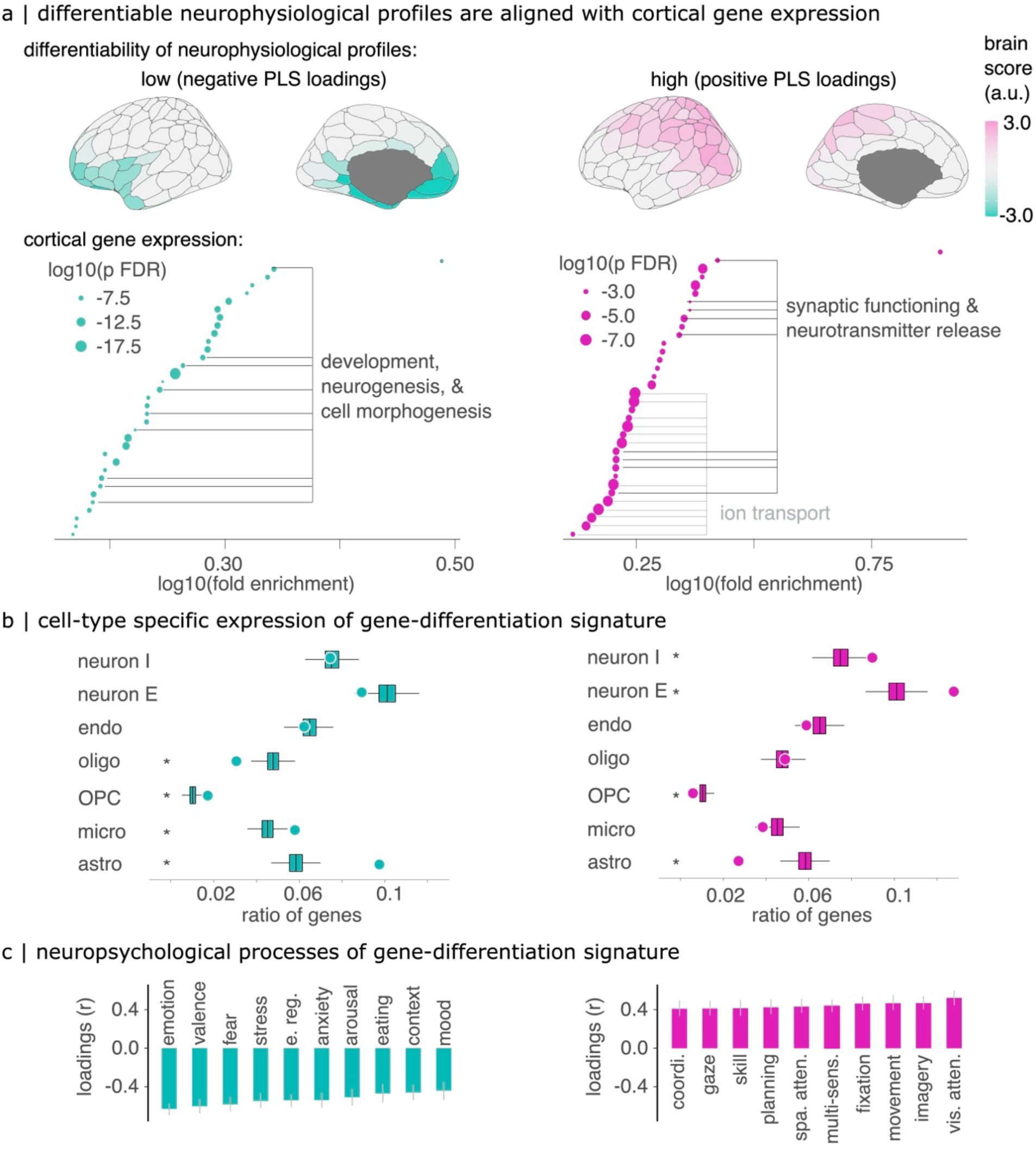
Associations Between Neurophysiological Traits, Cortical Gene Expression, and Neuropsychological Processes. (a) Gene-differentiation Partial Least Squares (PLS) analysis. The top panel shows neurophysiological brain score patterns for positive and negative loadings, indicating which cortical parcels align positively and negatively with the observed covariance pattern. The bottom panel presents the results of the gene ontology analysis. Each point represents an enriched biological process within the corresponding gene set. The size of each point denotes the associated p-value, while color intensity reflects the p-value after spatial autocorrelation correction (pSPIN). For clarity, related terms have been grouped together using horizontal bars. (b) Cell-type deconvolution analysis illustrating the proportion of genes (both positive and negative) preferentially expressed in seven distinct cell types, based on prior single-cell and single-nucleus RNA sequencing studies (*38–42*, *49*). The significance of these ratios was assessed via permutation testing (*p < 0.05*). Points represent observed ratios, while box plots show the distribution of permuted gene set ratios. Key: ‘neuron I’ refers to inhibitory neurons, ‘neuron E’ to excitatory neurons, ‘endo’ to endothelial cells, ‘oligo’ to oligodendrocytes, ‘OPC’ to oligodendrocyte precursor cells, ‘micro’ to microglia, and ‘astro’ to astrocytes. (c) Gene-Neuropsychological Processes PLS analysis. This panel illustrates the top 10 psychological terms that most significantly contribute negatively (cyan) and positively (pink) to the latent component identified in the gene-neuropsychological processes PLS analysis. The bar graphs display the loadings of neuropsychological terms. Confidence intervals were computed via bootstrapping and are not necessarily symmetric. Positively weighted terms positively covary with neurophysiological profiling and the positively weighted gene set.

Note that positively weighted frequency bands covary positively with positively weighted genes, and negatively relate to negatively weighted genes (see Methods). For example, cortical parcels with positive scores (Figure 3B) indicate covariance between positively weighted genes (Figure 3A) and positively weighted frequency bands, elucidating the relationship between cortical gene expression and participant differentiation.

To determine the types of cells corresponding to these genes, we analyzed gene sets that are preferentially expressed in seven cell types as determined by RNA sequencing studies(*38–43*). A ratio above the null-permuted values signifies that the given gene set is preferentially expressed in a specific cell type, while a ratio lower than the permuted values would indicate that the gene set is underexpressed in that cell type. We found an overrepresentation of genes with positive loadings in excitatory neurons (p_FDR_= 0.002; 1,000 gene permutations, two-tailed, FDR: corrected for false discovery rate) and inhibitory neurons (p_FDR_ = 0.006). Conversely, genes with positive loadings were underrepresented in astrocytes (p_FDR_ = 0.002) and oligodendrocyte precursor cells (OPCs; p_FDR_ = 0.03). Genes with negative loadings were predominantly represented in astrocytes (p_FDR_ = 0.002), microglia (p_FDR_ = 0.004), and OPCs (p_FDR_ = 0.002), but were less represented in oligodendrocytes (p_FDR_ = 0.002; Figure 3B left panel).

These findings suggest a clear dichotomy: genes that show positive loadings and thus positively correlate with the distinctive traits of neurophysiological profiles are predominantly expressed in neurons and underexpressed in OPCs and astrocytes. In contrast, genes with negative loadings, indicating a negative correlation with these neurophysiological profiles, are more frequently expressed in neuron-supporting cells such as astrocytes and microglia. Our results are consistent with existing models of the physiological origins of MEG signals (*27*, *44*, *45*).

### Association with Neuropsychological Processes

Building on prior work reporting associations between brain-fingerprint features and cognitive traits (*1–3*, *46*, *47*) as well as genetic influences on inter-individual variations in cognitive domains (*22–26*), we investigated how the gene-neurophysiological associations discovered in our data may relate to neuropsychological processes.

To investigate the relationship between neurophysiological differentiation and cognitive function, we utilized Neurosynth, a meta-analytic tool that aggregates fMRI activation data from over 15,000 studies (*48*). Neurosynth generates probabilistic cortical maps indicating the likelihood that activity in a given brain region is associated with a specific cognitive term (e.g., ’memory,’ ’attention’). Importantly, these maps do not distinguish between task-related activations and deactivations, instead reflecting the overall spatial distribution of reported cognitive associations.

To assess whether cortical gene expression aligns with cognitive function, we performed a Partial Least Squares (PLS) analysis comparing cortical expression patterns with Neurosynth-derived cognitive term maps. This allowed us to test whether the genes linked to neurophysiological differentiation also show spatial correspondence with cognitive processes.

A single significant component accounted for 67.2% of this covariance (CI [54.7, 72.2%], p_spin_ = 0.002), highlighting a distinction between cognitive and emotional domains (Figure 3C): negative loadings were associated with processes related to emotions, mood, and arousal, whereas positive loadings were linked to attentional, planning, and multimodal sensory processes.

We found that these patterns of covariance between gene expression and psychological processes were strongly aligned with those linking gene expression to individual neurophysiological traits (r=0.99, p_spin_ < 0.001; see also Supplemental Information). We interpret these findings to suggest that the identified pattern of gene expression covaries similarly with neurophysiological profiling and activations of psychological processes: the positive set of genes positively covaries with participant differentiation and processes such as attention and planning tasks, whereas negatively weighted genes covary with poor participant differentiation (i.e., low ICC) and processes including emotions and mood (Figure 3C).

### Developmental Trajectory of the Gene-Differentiation Signature

Previous work has established that genetic influences on neuropsychological processes become more pronounced with development (*24*, *26*, *50*). Following our identification of the cortical topography of a gene-differentiation signature that aligns with brain activations associated with neuropsychological processes (Figure 3A & C), we tested whether informative neurophysiological profiles in adulthood relate to cortical gradients of gene expression throughout development. To test this hypothesis, we assessed the topographical alignment between the cortical expression of genes across life stages in 12 cortical regions (*51*) and that of the gene-differentiation signature measured in adulthood. We found that this alignment (i.e., slope) was stronger in later life stages in all tested cortical regions, except for the hippocampus (Figure 4). These findings suggest that individual neurophysiological profiles in adulthood are most aligned to patterns of gene expression in adulthood, which become increasingly pronounced throughout development.

**Figure 4.**
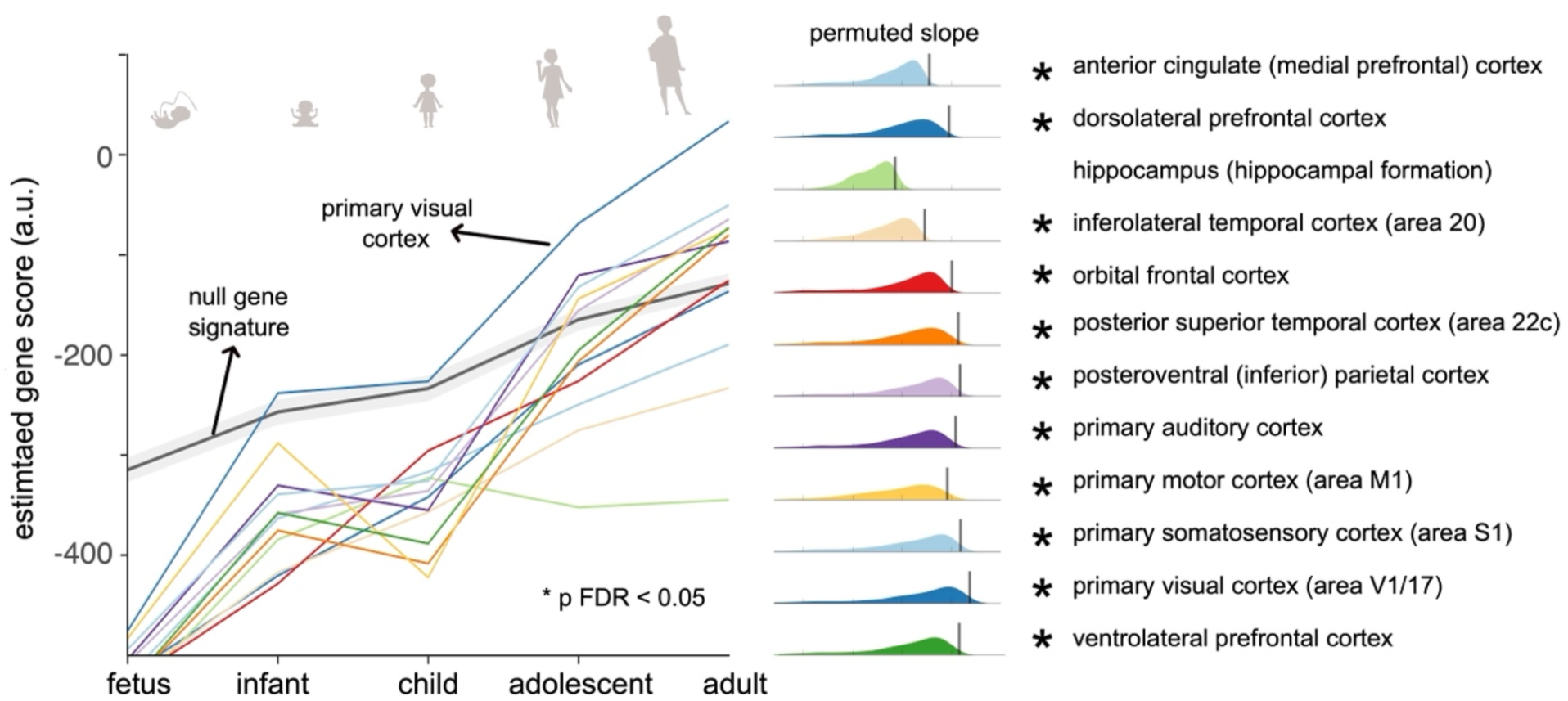
Strengthening of the Gene-Differentiation Signature Across Development. The left panel illustrates the progressive increase in gene-differentiation scores across developmental stages (prenatal through adulthood) for 12 brain regions, based on the BrainSpan data (*51*). A higher gene-differentiation score indicates greater similarity between the gene expression pattern at a given developmental stage and the gene-differentiation signature identified in adult neurophysiological profiles (see Figure 3). The solid black line with grey shading indicates the trajectory of gene scores derived from random permutations of gene expression data. The right panel shows histograms of permuted slopes for each cortical region; vertical lines represent the empirical slopes, and asterisks denote regions where the gene-differentiation signature significantly strengthens across development (*p_FDR_ < 0.05). Note that this analysis compares adult neurophysiological profiles with a longitudinal dataset of cortical gene expression. It therefore does not assume longitudinal changes in neurophysiological profiles, but rather assesses whether these adult profiles align with age-related changes in gene expression patterns.

## Discussion

Patterns of brain activity can differentiate between individuals, akin to hand fingerprints (*1–3*, *52*), and are associated with neuropsychological traits (*1*, *2*, *46*, *47*) and pathophysiology (6, 7, 30). We investigated the genetic bases of individual neurophysiological profiles. Our findings indicate that genetic factors explain a significant—but not complete—portion of the variance in neurophysiological profiles. These profiles covary with a cortical gradient of gene expression enriched for neurotransmission-related genes and preferentially expressed in areas supporting higher cognitive functions. This identified gene-differentiation signature becomes progressively pronounced throughout neurodevelopment, suggesting a growing genetic contribution to inter-individual variability in brain signalling.

### Heritability of Neurophysiological Traits

Our data show that some of the neurophysiological traits that shape individual profiles are heritable. The neurophysiological profiles of monozygotic twin pairs match each other beyond what can be explained by mere neuroanatomical resemblances (Figure 1B & Figure S3). Conversely, our ability to match the neurophysiological profiles of dizygotic twins was at random chance, achieving similar accuracy to noise recordings (Figure 1B). This suggests that the similarity between the neurophysiological profiles of dizygotic twin pairs is comparable to that of two unrelated individuals. These observations underscore that neurophysiological traits are in part shaped by genetics. In line with this interpretation, we observed that neurophysiological traits of alpha and beta-band activity were heritable (Figures 1, S5, & S6), confirming previous observations of the heritability of the individual frequency of alpha activity in humans (*16*, *17*).

The neurophysiological profiles most informative for participant differentiation followed similar spatial distributions to the heritable traits across alpha, beta, and gamma bands (Figure S4B & S6). These findings align with prior studies on MEG brain fingerprinting, which emphasize posterior sensorimotor regions—particularly in the beta band—as key for individual differentiation (*2*, *52*, *53*).

Our study demonstrates that genetic factors play a significant role in defining the uniqueness of neurophysiological profiles, yet they do not account for all observed variations. This discrepancy suggests an explanatory gap likely attributable to environmental influences. The inability to differentiate twin pairs with absolute precision underscores the need for further research with larger twin cohorts to fully elucidate the interplay between genetics and environment in shaping individual brain activity.

### A Gene-Differentiation Signature

Gene expression profoundly impacts brain structures and functions (*32*, *49*, *54–58*), such as cortical folding and connectivity within brain networks (*13*, *59*, *60*). Our study extends these findings by demonstrating that, in addition to anatomical and brain-network properties, genetics also shape task-free, ongoing neurophysiological brain activity. This activity reflects a set of distinctive neurophysiological traits, uniquely defining an individual’s profile in relation to the cortical expression of specific genes, forming a gene-differentiation signature.

Using gene expression atlases, we found that these traits are related to the expression of genes regulating ion transport and neurotransmission. Genetic variants and variations in gene expression levels likely influence the function of these genes, contributing to the observed inter-individual differences in neurophysiological profiles. This aligns with previous research that identified related genetic variants modulating alpha rhythms and other components of cortical neurodynamics (*16*, *18*, *61*). Our findings indicate that this gene signature is primarily active in both excitatory and inhibitory neurons (Figure 3B). Genetic influences on these cells likely manifest as observable differences in overall brain activity patterns, leading to distinct variations in macroscale brain signaling across individuals.

Conversely, our study revealed that the cortical expression of genes involved in cell morphogenesis and neurogenesis, particularly in limbic regions, exhibits a negative correlation with individual neurophysiological differentiation (i.e., different signed loadings in the PLS analysis; see Figure 3B). In contrast to positively-loaded genes, which relate to neurochemical interactions between neurons, these negatively-loaded genes are mainly expressed in supportive cells such as astrocytes and microglia. This suggests that genetic influences on neurophysiological individual traits are more related to neuronal communication than to inherited structural brain features, consistent with existing models of the physiological origins of MEG signals (*27*, *44*, *45*).

Our research highlights potential pathways for future studies on individual neurophysiology by providing a biologically grounded framework to understand behavioural variations. Animal models could be instrumental in manipulating alleles of the genes identified in our study to assess their impact on gene product functioning, large-scale brain signal characteristics, and ultimately, behaviour. This approach would provide further insights into how genetic variations influence both neurophysiological traits and behavioural differences.

### Alignment of the Gene-Differentiation Signature with Neuropsychological Processes

We found that, across the cortex, the gene-differentiation signature of neurophysiological profiles aligns with maps of cortical activations related to specific cognitive and emotional processes (Figure 3C). The spatial alignment between cortical gene expression and Neurosynth-derived cognitive maps reflects an organization pattern consistent with previously reported functional gradients (*62–65*). Specifically, the observed differentiation aligns with a cognitive-affective gradient, where sensory-driven cognitive processes (e.g., attention, perception) are spatially distinct from affective and emotion-related processes. These results suggest that individual differences in neurophysiology are embedded within this large-scale functional architecture.

The cortical gradients identified herein—including inter-individual neurophysiological differentiation, gene expression patterns, and cognitive functional associations—closely resemble the well-documented unimodal-to-transmodal axis of cortical organization (*62–65*). This large-scale gradient is a fundamental organizing principle of the brain, observed across multiple levels of neurobiology, including structural connectivity, functional networks, and transcriptomic variation (*63*, *65*)

Our findings suggest that inter-individual variability in neurophysiological traits follows this primary axis of cortical organization. Sensory areas, which exhibit the highest neurophysiological differentiation, are functionally and molecularly distinct from transmodal association areas. These results extend prior literature by demonstrating that this organizational framework also applies to individual differences in large-scale neurophysiology. Future studies should explore the genetic and developmental factors shaping these gradients and their role in cognitive and behavioral variability.

### Trajectory of the Gene-Differentiation Signature Throughout Development

Prior research has shown that genetic influences on neuropsychological processes increase across development (*24*, *26*, *50*) and that individual brain-fingerprint features become more stable and unique with age (*5*, *66*). To examine whether neurophysiological differentiation aligns with gene expression across the lifespan, we assessed the spatial alignment between adult neurophysiological differentiation maps and cortical gene expression gradients in 12 cortical regions (51). We found that this alignment (i.e., slope) was strongest in later developmental stages in all regions except for the hippocampus (Figure 3).

These findings suggest that the molecular landscape supporting neurophysiological differentiation in adulthood becomes increasingly structured throughout development and provide a preliminary molecular framework for understanding age-related changes in neurophysiological variability (*67–72*). Our findings are supported by independent evidence demonstrating increasing genetic influences on cognition and brain activity across the lifespan (*24*, *26*, *50*). Additionally, a recent complementary study on a lifespan dataset (>1000 individuals, ages 4–89) suggests a progressive shift from sensory to transmodal regions in individual differentiation, paralleling the developmental trajectory of gene expression gradients (*53*).

### Methodological Considerations and Future Directions

Previous neuroimaging studies with fMRI have shown how gene expression modulates functional connectivity in the frontoparietal network, a key feature of inter-individual differentiation in fMRI brain-fingerprints (*73*). In contrast, our study highlights the role of posterior unimodal sensory cortical regions in driving inter-individual differentiation based on neurophysiological traits. This disparity between neuroimaging modalities underscores the distinct biological underpinnings of hemodynamic fMRI and electrophysiological signals (*1*, *2*, *52*, *74*). Specifically, while fMRI connectome-based individual profiles partially reflect the structural connections between network nodes (*75*, *76*), our findings indicate that individual neurophysiological profiles are predominantly shaped by processes of neuronal communication, particularly through the expression of genes involved in ion transport and neurotransmission. This distinction underscores the critical role of these genes in mediating neuronal signaling and brain activity patterns, providing novel insight into the mechanisms underlying neurophysiological individuality.

We must also underscore potential limitations in the interpretation of our findings. Heritability estimates were derived using Falconer’s formula, which does not account for variance due to environmental factors. Moreover, due to our limited sample size, we do not interpret heritability estimates in terms of absolute variance explained by genetics alone. Instead, we emphasize the spatial distribution of heritable neurophysiological traits and their alignment with neurophysiological differentiation patterns (Figures S4 & S6). Future studies with larger twin cohorts and comprehensive demographic documentation are warranted to refine these estimates further. Our genomics analysis is based on rare, albeit limited, data from a small sample of post-mortem brain tissue. The tissue sampling process has inherent biases, such as a focus on the left hemisphere and sex imbalance. Future research should aim to mitigate these biases. Additionally, we acknowledge that post-mortem gene expression measures may not accurately reflect in vivo conditions (*77*).

A specific strength and weakness of our approach lies in aggregating data from multiple sources. While this integration uniquely enables addressing multiscale neuroscientific questions, it also presents inherent methodological challenges. To mitigate concerns regarding data aggregation (e.g., alignment between cortical maps from the Allen Human Brain Atlas and the Human Connectome Project), we cross-validated our PLS results on a held-out sample of cortical parcels (Figure 2C) and employed spatial autocorrelation-preserving permutations. Our findings do not imply, however, that individual deviations in gene expression predict deviations in neurophysiology, nor do they imply direct longitudinal tracking of gene expression within individuals in the case of the BrainSpan dataset. Instead, they suggest that neurophysiological differentiation follows an organizational pattern that aligns with normative cortical gene expression. The correlational nature of our findings highlights the need for further experimental validation, potentially through animal models, to establish causative links between gene expression and neurophysiological and behavioural traits.

In conclusion, our research elucidates the relationship between molecular variations, brain activity, and individual differences. Using a multiscale, data-driven approach, the present study suggests new avenues for understanding the biological foundations of individual variability. Our findings lay the groundwork for future studies to further explore these complex interconnections, thereby enriching our understanding of the neural underpinnings of human behaviour and cognition. We hope that our work will inspire continued exploration and innovation in the field, ultimately advancing our knowledge of how genetic and neurophysiological factors shape the human experience.

## Methods

### Participants

MRI and MEG data from 89 healthy young adults (22-35 years old; mean= 28.6, SD= 3.8 years; see Table S1) were collected from the Human Connectome Project (HCP)(*29*). Among these 89 participants, 34 were monozygotic twins, and 22 were dizygotic twins. The zygosity of the participants was confirmed with genotyping tests. All participants underwent three approximately six-minute resting-state eyes-open MEG recordings using a 248-magnetometer whole-head Magnes 3600 system (4DNeuroimaging, San Diego, CA). All sessions were conducted at the same location with a sampling rate of 2034.5 Hz, as detailed in HCP protocols(*29*). Data from twin pairs included in the Human Connectome Project were sex-matched and reared similarly (*29*, *78*). This ensures that observed differences in neurophysiological differentiation between MZ and DZ twins are not attributable to differences in early-life environmental exposure.

### Ethics

The procedures for the curation and analysis were reviewed and approved according to the institutional ethics policies of McGill University ’s and the Montreal Neurological Institute’s Research Ethics Boards (ref no. 22-06-079).

### MEG Data Preprocessing & Source Mapping

EG data were preprocessed following good practice guidelines(*79*) using Brainstorm(*80*). Source maps for each participant’s recordings were computed using a linearly-constrained minimum-variance (LCMV) beamformer and were clustered into 200 cortical parcels of the Schaefer atlas (*36*), as detailed in Supplemental Information Methods.

### Neurophysiological Profiles

Power spectrum density (PSD) estimates at each cortical parcel were derived using Welch’s method (sliding window of 2 s, 50% overlap). The neurophysiological profile (or *brain-fingerprint*) of each participant consisted of PSD values defined at 301 frequency bins (range: 0-150Hz; ½ Hz resolution) for each of the 200 cortical parcels. Neurophysiological profiles were generated for each of the three MEG recordings per participant.

### Individual Differentiation

Individual neurophysiological profiling was conducted following our previous work (Figure 1A)(*2*). We assessed the correlational similarity between participants’ neurophysiological profiles across recordings. For each probe participant, we computed Pearson’s correlation coefficients between their neurophysiological profile from one their three recordings available and a test set consisting of the neurophysiological profiles of all participants derived from another one of the other two recordings (between-participants similarity), including the probe participant’s profile (within-participant similarity). A participant was correctly differentiated if the highest correlation coefficient between their neurophysiological profile and the test set was obtained from their own neurophysiological profile from the other recording. This procedure was repeated for all participants. We then computed the percentage of correctly differentiated participants across the cohort, yielding a score of differentiation accuracy for the neurophysiological profiling approach. This procedure was repeated for all possible pairs of data recordings from the three available for each participant, and the mean differentiation accuracy was reported.

### Matching Neurophysiological Profiles Between Twin Pairs

We declared that the neurophysiological profiles of twin siblings matched with one another if their Pearson’s correlation coefficients were higher than with any other participant. This matching procedure was repeated for all twin pairs in the cohort, and we reported the percentage of correctly matched pairs separately for monozygotic and dizygotic twin pairs. The results of this analysis are presented in Figure 1b.

To test chance-level matching between neurophysiological profiles, we assessed our ability to match the neurophysiological profiles of randomly paired unrelated individuals. We randomly assigned the unrelated individuals in the cohort to another person’s neurophysiological profile and computed the matching accuracy. We repeated the randomization procedure 300 times for each frequency band.

### Band-limited Neurophysiological Profiles

We replicated the individual and twin pair neurophysiological profiling analyses, restricting the PSD features to those averaged over the typical electrophysiological frequency bands: delta (1–4 Hz), theta (4–8 Hz), alpha (8–13 Hz), beta (13–30 Hz), gamma (30–50 Hz), and high-gamma (50–150 Hz). For example, the alpha-band neurophysiological profile consists of power spectral density estimates at each frequency within 8–13 Hz, rather than an average across this range.

### Bootstrapping of Differentiation Accuracy Scores

To derive confidence intervals for the reported differentiation accuracies, we employed a bootstrapping method. We randomly selected a subset of participants representing approximately 90% of the tested cohort (i.e., 30 MZ, 20 DZ, and 30 non-twins), derived a differentiation accuracy score, and repeated this procedure 1000 times with random subsamples of participants. We report 95% confidence intervals from the 2.5^th^ and 97.5^th^ percentiles of the resulting empirical distribution of differentiation accuracies. For deriving confidence intervals for the differentiation of twin pairs, we randomly subsampled 15 MZ twin pairs and 10 DZ twin pairs for each iteration of the random subsampling.

### Saliency of Neurophysiological Traits

We calculated intraclass correlations (ICC) using a one-way random effects model to quantify the contribution of each cortical parcel and frequency band toward differentiating between individuals across the cohort (*3*, *81*). This approach follows prior work introducing ICC for characterizing topological fingerprints in neuroimaging (*3*, *81*). ICC quantifies the ratio of within-participant to between-participant variance. High ICC values indicate the saliency of a feature of the neurophysiological profile (a neurophysiological *trait*) in distinguishing individuals, as it reflects high within-participant consistency and low between-participant variability. To avoid potential bias due to twin pairs, we computed ICC across all individuals in the cohort and across 100 random subsamples, ensuring only one twin from each pair was included in each subsample (i.e., for each subsample, we randomly selected either twin A or twin B to include in the calculation of ICC). The ICC values obtained from bootstrapping were nearly identical to those obtained from the entire cohort (98.6% correlation). We proceeded with the ICC values averaged across bootstraps for all analyses. The results of this analysis are presented in Figure S4b.

Chance-level participant differentiation: To rule out the possibility that environmental factors influenced participant differentiation and twin matching accuracies, we computed a measure of chance-level differentiation and matching accuracy using mock neurophysiological profiles derived from empty-room noise recordings collected during each MEG session. These recordings, obtained from the HCP dataset (*29*), capture instrumental and environmental noise such as: Time of day effects (e.g., variations in background electromagnetic activity); scanner-related variability (e.g., fluctuations in baseline signal sensitivity); external noise sources (e.g., minor variations in shielding effectiveness or room conditions).

From these empty-room noise recordings, we derived a single ’mock brain-fingerprint’ for each participant, based on our previous work (*2*, *8*, *53*). The noise time series was projected onto each participant’s cortical source maps, and differentiation accuracy was recalculated using these mock profiles to test whether chance-level accuracy varied systematically across frequency bands. These were projected onto participant cortical source maps, and differentiation was recalculated using the same pipeline. Importantly, chance-level differentiation remained low (<15%) across all frequency bands, and the alpha band did not show disproportionate differentiation accuracy, ruling out SNR-driven bias.

### Heritability of Neurophysiological Traits

We calculated the heritability of individual neurophysiological profiles, considered as phenotypes, using the Falconer formula (*11*). This method estimates the relative contribution of genetics versus environmental factors in determining a phenotype. A phenotype in this context refers to the overall neurophysiological profile of an individual, while a trait refers to specific aspects or features within this profile, such as power spectrum density in a particular frequency band or cortical region. If the similarity in a phenotype between monozygotic (MZ) twins is greater than that between dizygotic (DZ) twins, the trait is considered heritable:

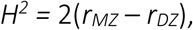

where *r_MZ_* (and *r_DZ_*, respectivey) is the intraclass correlation between MZ (and DZ, respectively) twin pairs for a given neurophysiological trait. It is important to note that heritability reflects the similarity within twin pairs for a given phenotype, whereas ICC reflects the stability of a trait within a person relative to others in the cohort.

Heritability estimates were computed using three recordings per twin, resulting in 9 pairwise comparisons per twin pair (i.e., 3 × 3 comparisons) following a cross-twin cross-session pairwise comparison.

To confirm the robustness of our results, we tested whether heritability remained stable across different sample sizes and correlation metrics. Subsampling MZ twins to match DZ sample sizes resulted in highly similar heritability estimates (r = 0.90). Additionally, using Pearson’s correlation instead of ICC produced near-identical estimates (r = 0.98).

### Gene Expression Data

Gene expression data was obtained from the six postmortem brains provided by the Allen Human Brain Atlas (AHBA; http://human.brain-map.org/) (33) using the *abagen* python package(*35*). Our analyses followed a similar pipeline to prior studies(*32*). Gene expression was obtained by averaging across donors. We computed differential stability [Δ_S_(p)] for every gene probe. Differential stability is defined as the consistency of gene expression patterns across different donor brains, quantified using Spearman rank correlation. Higher differential stability indicates that a gene exhibits spatially stable expression patterns across individuals, making it more suitable for population-level analyses. We retained 9104 genes with a differential stability above 0.1 for all future analyses, following good-practice guidelines and previous literature(*32*, *34*, *35*, *82*). See Supplemental Information Methods for further details.

### PLS Derivation of a Gene-Differentiation Signature

We related features for participant differentiation to gene expression gradients using partial least squares (PLS) analysis (*83–85*). We z-scored the columns of two data arrays: one containing the full neurophysiological trait matrix (ICC values across frequency bands) and the other containing gene expression levels. The neurophysiological traits array (denoted as *Y*) had 6 columns representing each frequency band of interest and 200 rows representing the cortical parcels of the Schaefer atlas. The gene expression array (denoted as *X*) had 9104 columns representing genes and 200 rows representing the same cortical parcels. We applied singular value decomposition (SVD) to the covariation matrix of *X* and *Y* such that:

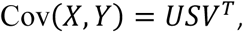

where *U* and *V* are the left and right singular vectors, and *S* is the diagonal matrix of singular values. As typical with PLS, this decomposition allowed us to identify the latent variables that maximally covary between the gene expression data and the neurophysiological traits. Each singular value indicates the amount of covariance explained by its corresponding latent component. The vectors *U* and *V* provide the weights for the genes and frequency bands, respectively, for each latent component. High weights in *U* correspond to genes that strongly covary with high weights in *V*, which correspond to specific frequency bands. Positively weighted genes covary with positively weighted frequency bands, elucidating the relationship between genetic and neurophysiological variability.

Gene expression and ICC scores were computed for each cortical parcel by projecting the original data matrices *X* and *Y* onto the singular vector weights obtained from the PLS analysis. Specifically, these scores represent the covariance between the gene expression data and the neurophysiological traits. For example, cortical parcels with positive scores indicate covariance between positively weighted genes and positively weighted frequency bands, which are important for participant differentiation.

The contribution of each frequency band to the observed latent variable (i.e., loadings) was computed using Pearson correlation between ICC values and the cortical score pattern obtained from PLS analysis. These correlations represent the loadings for each frequency band, quantifying their alignment with the gene-differentiation signature. Loadings were computed as Pearson’s correlation coefficients between each variable’s regional spatial distribution over the cortex (i.e., gene expression and ICC data) and the corresponding cortical score pattern (i.e., correlating gene expression with ICC scores). We used Pearson’s correlation coefficients for our loadings because they provide a standardized measure of the strength and direction of the linear relationship between variables, facilitating interpretation. Variables with large absolute loadings are highly correlated with the observed score pattern and strongly relate to the latent component of covariance.

To assess the significance of the latent components, we conducted permutation tests that preserved the spatial autocorrelation of cortical maps (see below). Specifically, we performed 1,000 spin tests and computed a null distribution of singular values. P-values were computed as the proportion of null singular values that achieved a greater magnitude than the empirical singular values. Additionally, we computed bootstrapped confidence intervals for the singular values by randomly resampling the rows (corresponding to the cortical parcels) of both data matrices 1,000 times. We report the 2.5^th^ and 97.5^th^ percentiles of the resulting distribution of singular values. The results of this analysis are presented in Figure 3a.

### Gene Ontology Analysis

To determine the biological processes that strongly contributed to the set of positively and negatively loaded genes, we conducted an enrichment analysis using gene ontology(*86*, *87*), a framework for categorizing gene products based on their molecular function and associated biological processes(*37*, *86*).

For negatively and positively loaded genes, we separately selected the 50% with the largest absolute loadings (e.g., genes with the 50% most negative or positive loadings) and input these genes into the *ShinyGO v.0.77* gene ontology tool (*37*) using the GO biological processes pathway databases(*87*). *ShinyGO* is an accessible bioinformatics Shiny app which conducts gene ontology analyses to identify which biological processes are enriched in any given set of genes. Genes without Enrtez IDs were excluded from the analysis. P-values associated with fold enrichment for all terms were FDR corrected. See Supplemental Data for a comprehensive list of all biological processes and their corresponding fold enrichment values. The results of this analysis are presented in Figure 3c.

Previous research has shown that standard gene ontology analyses may inflate significance estimates due to spatial autocorrelation in cortical maps (*88*). To mitigate this issue, we employed spatial autocorrelation-preserving permutation tests to rigorously evaluate statistical significance (see Methods: Correction for Spatial Autocorrelation of Cortical Maps). Specifically, for each gene ontology category, we computed the spatial alignment between gene expression topographies and neurophysiological differentiation maps, then assessed significance against 1000 iterations of spatially permuted cortical maps. This approach ensures that our findings exceed what would be expected under a random spatially autocorrelated null model.

Critically, all previously reported gene ontology categories remained significant following these rigorous control analyses, confirming the robustness of our findings. These results are now illustrated in Figure 3A (see pSPIN values).

### Cell-type Deconvolution

We aggregated cell-specific gene sets for seven cell types using data from five human adult postmortem single-cell and single-nucleus RNA sequencing studies(*38–42*, *49*). The seven cell classes were determined based on hierarchical clustering, resulting in the following cell types: astrocytes, endothelial cells, microglia, excitatory neurons, inhibitory neurons, oligodendrocytes, and oligodendrocyte precursor cells (OPC).

We assessed the preferential expression of cell-specific gene sets by 1) computing the ratio of positively loaded genes that overlapped with the cell-specific gene set; and 2) permuting gene sets 1,000 times to assess statistical significance.

This approach allowed us to determine the statistical significance of the overlap between the loaded genes and the cell-specific gene sets, providing insights into the cell-type-specific expression patterns of the genes contributing to the neurophysiological traits. The results of this analysis are presented in Figure 3b.

### Gene Expression & Neuropsychological Processes PLS Analysis

We repeated the above-described PLS analysis to relate gene expression to neuropsychological processes, as indexed by brain activation maps obtained from *Neurosynth*(*48*). This analysis replicated the approach of Hansen and colleagues(*32*), using the Schaefer-200 atlas(*36*). For detailed methodology, see Supplemental Information Methods.

### Gene-Signature Evolution Across Developmental Stages

We used brain gene expression data available from BrainSpan(*51*) ,which features gene expression levels from different developmental stages ranging from 8 post-conception weeks to 40 years of age. We computed gene scores for 12 cortical regions across neurodevelopmental stages by multiplying the gene expression matrix obtained from BrainSpan with the PLS-derived gene weights (columns of *U*) obtained from the gene-differentiation PLS analysis (see **PLS Derivation of a Gene-Differentiation Signature**).

We fitted linear slopes—using the MATLAB *polyfit()* function—to the gene scores across neurodevelopment for each cortical parcel separately. These slopes were then compared to statistical null slope values obtained by performing spatially autocorrelation-preserving permutations, then running the PLS analysis pipeline, and multiplying the null gene weights with BrainSpan gene expression data. This resulted in a null distribution of slopes (1,000 permutations). See Supplemental Information Methods for further details. The results of this analysis are presented in Figure 3 and S7.

### Visualization

We plotted brain maps of ICC, heritability, and PLS brain scores using the *ggSchaefer* and *ggseg* R packages. All other plots were generated using the *ggplot2* package in R(*89*).

### Correction for Spatial Autocorrelation of Cortical Maps

We corrected for spatial autocorrelation of cortical map data, where applicable, using SPIN tests. SPIN tests preserve the spatial autocorrelation of cortical topographies by rotating the cortex surface data, effectively permuting the spatial positions while maintaining the spatial structure. This generates a null distribution that accounts for spatial autocorrelation. We conducted 1,000 spin permutations of our brain maps using the Hungarian method(*90*, *91*).

## Acknowledgements

The funders had no role in study design, data collection and analysis, decision to publish, or preparation of the manuscript. The Brainstorm app is supported by funding to SB from the NIH (R01-EB026299), a Discovery grant from the Natural Science and Engineering Research Council of Canada (436355-13), the CIHR Canada Research Chair in Neural Dynamics of Brain Systems, the Brain Canada Foundation with support from Health Canada, and the Innovative Ideas program from the Canada First Research Excellence Fund, awarded to McGill University for the HBHL initiative. This work was supported by a doctoral fellowship from NSERC (JDSC, JYH).

## Author Contribution

Conceptualization: JDSC, SB

Data Curation: JDSC, SB

Methodology: JDSC, JP, JYH, BM, SB

Software: JDSC, JP, JYH, SB

Visualization: JDSC, JP, SB

Resources: JYH, SB

Validation: JDSC, SB

Formal analysis: JDSC

Supervision: SB

Project administration: SB

Funding acquisition: SB

Writing – original draft: JDSC, SB

Writing – review & editing: JDSC, JP, JYH, BM, SB

## Data and materials availability

All data needed to evaluate the conclusions in the paper are present in the paper and/or the Supplementary Materials. The data are available through the Human Connectome Project (HCP) repository (https://www.humanconnectome.org/study/hcp-young-adult). Gene expression data are available through the Allan Human Brain atlas (ABHA; http://human.brain-map.org/). The Neurosynth database is available at https://neurosynth.org/.

All in-house code used for data analysis and visualization is available on Zenodo https://zenodo.org/records/15023999 and our GitHub https://github.com/Epideixx/Fingerprints_Twins.

## Supplemental Materials

### Methods (Cont’d.)

#### MEG data preprocessing

MEG data were preprocessed following good practice guidelines (*1*) using *Brainstorm* (*2*) March-2023 distribution running MATLAB 2020b (Mathworks Inc., Massachusetts, USA). Our preprocessing pipeline was adapted from previous published work(*3*, *4*). Line noise artifacts (60Hz) along with their first 9 harmonics were removed using notch filters. Slow-wave and DC artifacts were attenuated with a high-pass FIR filter above 0.3 Hz. To remove ocular and cardiac physiological artifacts, we defined Signal-Space Projections (SSPs) based on the activity of concurrent electro-cardiogram and -oculogram recordings. We additionally attenuated low-frequency eye saccades (1-7 Hz) and high-frequency (40-240 Hz) muscle noise components with SSPs.

#### MEG source mapping

We source imaged the resting-state MEG sensor data using the coregistered anatomy folder provided by HCP(*5*). We computed MEG biophysical head models for each participant using the *Brainstorm* overlapping-spheres model (default parameters) applied to 15,000 locations distributed over the entire cortex. Source maps for each participants’ recording were computed using linearly-constrained minimum-variance (LCMV) beamforming (using *Brainstorm*‘s default parameters: 2018 version). Noise statistics were estimated from the empty-room recordings collected on the respective day of the visit of each participant. Individual source maps were then projected onto a default anatomy template, spatially smoothed (3mm) and clustered into the 200 cortical regions of the Schaefer atlas(*6*) using the first principal component within each region as a representative time series of brain activity. Brain-fingerprints were derived from the power spectrum densities (PSD) of these regional source time series computed using Welch’s method with a sliding window of 2 seconds and 50% overlap.

### Correspondence of salient neurophysiological traits and heritable brain phenotypes

We determined whether the salient features for individual differentiation were aligned topographically with heritable brain phenotypes. To do this, we computed the Pearson’s spatial correlation of ICC neurophysiological profile topographies with the brain maps obtained from the heritability analyses (see **Heritability of brain phenotypes**) across the 200 regions of the Schaefer atlas(*6*). We controlled for the spatial autocorrelation of the data using the Hungarian method (*7*, *8*) (see **Correction for spatial autocorrelation of brain maps**).

### Neuroanatomy

We verified that the neurophysiological profiles of MZ twin pairs matched in spite of heritable neuroanatomical features. We, therefore, derived structural statistics for each region of the Desikan-Killiany atlas from *Freesurfer* (*9*). We then i) computed the heritability of these features following the procedure described in the main text, and ii) tested for a possible linear association between anatomical and spectral similarity across twin pairs. The results are reported separately for MZ, and DZ twin pairs (see The Matching Between the Neurophysiological Profiles of Monozygotic Twins Is Not Driven by Anatomy).

### Biophysical and environmental artifacts

We investigated whether MEG recording artifacts might have overly contributed to the differentiation between individuals. We computed the root-mean-squares (RMS) of ocular and cardiac reference signals (ECG, HEOG, VEOG, respectively) collected simultaneously with MEG data. We then linearly regressed these measures from the neurophysiological profiles and used the residuals of this regression to differentiate individuals. We then tested whether the environmental and instrument noise conditions on the day of the MEG recordings biased individual differentiation(*3*). We, therefore, used the empty-room recordings collected on the same day of the MEG session for each participant to derive pseudo-neurophysiological profiles. These empty-room recordings were preprocessed using the same filters as the resting-state data and projected onto the participant’s brain using the same imaging kernels. We computed the differentiation accuracies obtained based on these pseudo-profiles.

### Gene expression data

Gene expression data were obtained from the six postmortem brains provided by the AHBA (http://human.brain-map.org/)(10) using the *abagen* Python package(*11*), following a pipeline published previously(*12*). In brief, we first used microarray probes with the highest differential stability to represent gene expression for each gene (20,232 in total). Tissue samples were assigned to each of the 200 brain regions of the Schaefer atlas using Montreal Neurological Institute (MNI) coordinates generated via nonlinear registrations. We ignored tissue samples further than 2 mm away from each brain region. To reduce potential misassignment, sample-to-region matching was constrained by hemisphere and to the cortex. If a region of the Schaefer atlas was not assigned a sample, the closest sample in Euclidian distance to the centroid of the region was selected. Gene expression was normalized across tissue samples and subjects, and for each of the retained genes, was obtained by averaging across donors. We retained 9104 genes with a differential stability above 0.1 in further analyses, following good-practice guidelines and previous literature(*11–14*).

### Cross-validation of gene-differentiation PLS analysis

We assessed the robustness of our PLS model through cross-validation of Pearson’s correlation between the observed gene scores and ICC statistics. We followed the same cross-validation procedures as Hansen and colleagues(*12*), splitting brain regions into one testing and one training set. A random seed was used to determine the training set: 75% of the brain regions the closest in Euclidian distance to the seed location were used to train the PLS model. The quartile of regions were held out to test the PLS model by computing the correlation between predicted gene scores and ICC statistics [*Corr(XtestUtrain, YtestVtrain*)]. This procedure was repeated 100 times to produce a distribution of correlations. The significance of the cross-validation outcomes was assessed against a null model obtained from spatial autocorrelation-preserving permutations of the gene expression matrix and repeated the cross-validation procedure 1,000 times (Figure S7c).

### Gene expression & psychological-processes PLS analysis

We assessed the relationship between gene expression and psychological processes as indexed by brain activation maps obtained from *Neurosynth*(*15*).

The brain map associated with each psychological-process term represents the probabilistic association between this term (e.g., attention) and brain activations observed at each voxel from published studies reporting on that psychological process. This meta-analytic approach combines data from >14,000 published fMRI studies. We focused our analyses on the 123 terms reported by Hansen and colleagues(*12*) at the intersection between Neurosynth(*15*) and the Cognitive Atlas(*16*), a public ontology of cognitive science. This data-driven approach did not distinguish between activations and deactivations, nor did it consider the degree of activation of a given brain area. Here too, we used the Schaefer-200 atlas(*6*) to sample the resulting cortical maps.

We assessed the alignment between the respective latent components associated with gene-psychological processes and gene-differentiation by computing the Pearson’s correlation between gene scores and PLS loadings (see Psychological Processes and Differentiation).

### Gene ontology analysis

To determine the biological processes contributing to positively and negatively loaded genes, we performed an enrichment analysis for the 50% largest loadings (e.g., genes with the 50% most negative and positive loadings) using the *ShinyGO* V 0.77 (Dec 20^th^ 2023) gene ontology tool(*17*) and the GO pathway databases of biological processes(*18*). Genes with no Entrez Ids were ignored. Fold enrichment for each biological process was computed by comparing the frequency of a given biological process in the set of positive genes to the frequency of that process in the entire genome. P-values associated with fold enrichment for all terms were corrected for false discovery rate (FDR). See Supplemental Data for a comprehensive list of all biological processes and their corresponding fold enrichment values.

### Development of the gene signature

We binned gene expression data from BrainSpan (*19*) into five life stages: fetal (8–37 post-conception weeks), infant (4 months–1 year), child (2–8 years), adolescent (11–19 years) and adult (21–40 years)(*20*). For each life stage, we computed the gene expression of the top 50% of positively and negatively loaded genes for each cortical region. Additionally, we computed gene expression at every neurodevelopmental stage for a random set of genes (Figure S7B). Note that of the 16 cortical regions with gene expression data, four regions only had samples for the fetal stage; therefore, we report data for the 12 cortical regions with data across all neurodevelopmental stages.

### Human accelerated region analyses

We defined genes associated with human accelerated regions (HARs) based on previous work by Wei et al.(*21*). Of the 1711 genes featured in AHBA, they reported that 415 genes were significantly more expressed in brain tissues than other available body samples (see Supplementary Data 2 therein). We used these 415 HAR-brain genes for further analyses, including 313 genes that were differential stable in our analysis (see **Gene expression data**). We first assessed the overrepresentation of HAR-brain genes in the identified gene signature (top 50% positive and negative loadings) through permutation analyses. We then assessed the spatial correspondence of gene expression of HAR-brain genes and the computed gene brain score (see **Correction for spatial autocorrelation of brain maps**).

## Supplementary Results

**Table S1:** Demographic information.

| | Age (mean ± SD) | Sex (F/M) | Education (mean ± SD) |
| --- | --- | --- | --- |
| Non-twins) | 29.08 ± 3.30 | 13/12 | 13.8 ± 2.16 |
| Monozygotic | 27.45 ± 3.94 | 16/22 | 15.31 ± 1.34 |
| Dizygotic | 29.85 ± 3.87 | 12/14 | 15.00 ± 1.57 |

### Assessing the Robustness of Neurophysiological Profiles

We performed a series of sensitivity analyses to rule out the possibility of environmental and physiological artifacts affecting our results.

We first evaluated the influence of environmental factors that may affected the MEG recordings. We processed empty-room recordings in the same way as the actual participant data to derive pseudo-neurophysiological profiles related to the environmental conditions around each participant’s visit. Individual differentiation was poor based on these pseudo-neurophysiological profiles (<1.7%; Figures 1B).

We then used linear regression models to remove the variance associated with physiological artifacts from the neurophysiological profiles (see Methods (Cont’d.) **Biophysical and environmental artifacts**). Using the same analysis pipeline, we observed that identification accuracy remained largely unaffected: 82.6% [74.7, 89.3] differentiation accuracy across all participants, 55.2% [46.7, 66.7] matching accuracy between monozygotic twins, and 5.8% [0.0, 15.0] between dizygotic twins, using broadband features (1-150Hz; Figure S2). The robustness of the results indicates that individual differentiation is not significantly driven by physiological artefacts.

**Figure S1.**
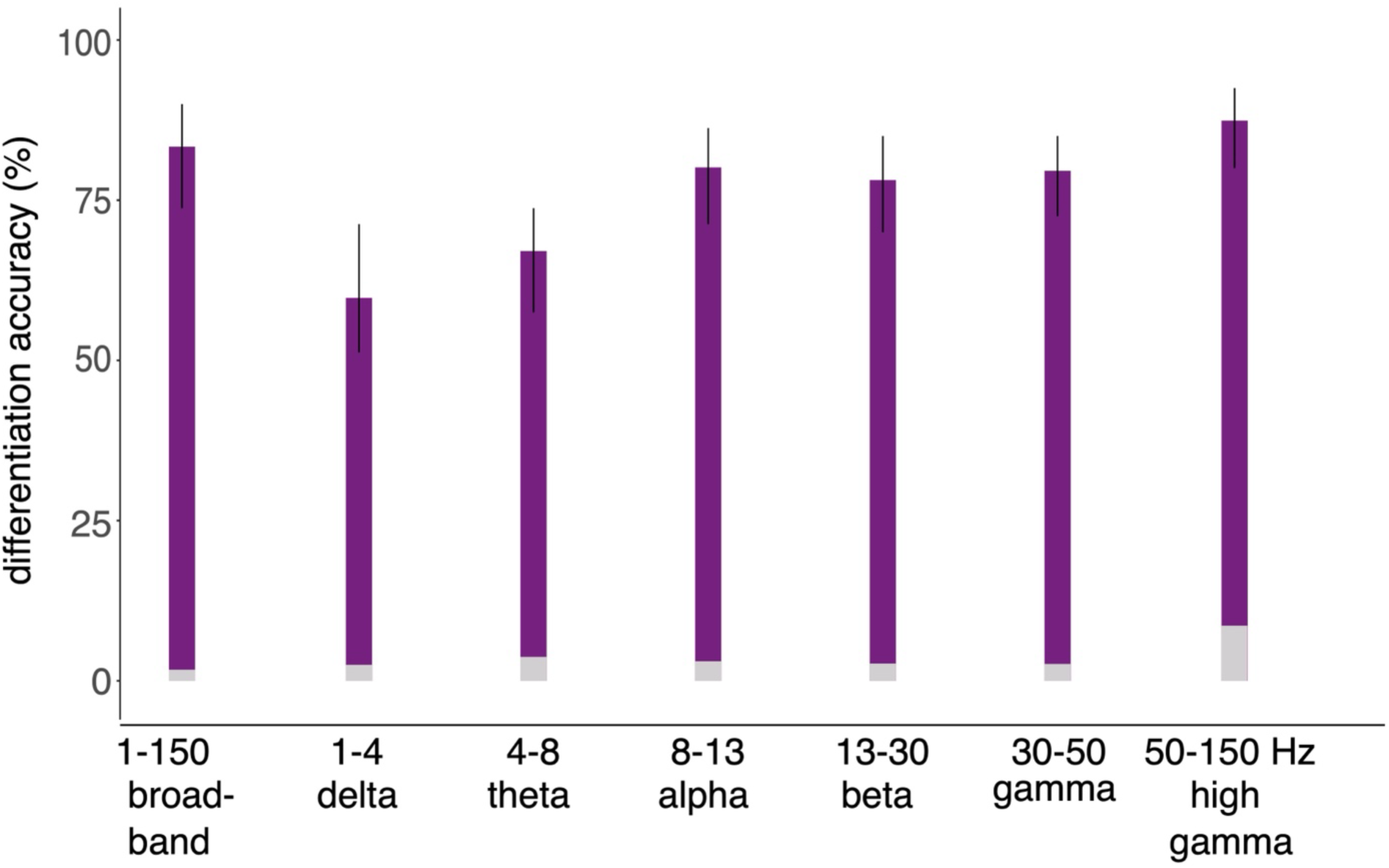
Participant differentiation accuracy. The differentiation accuracy scores of neurophysiological profiles for all individuals. Participants can be accurately differentiated from neurophysiological profiles across all frequency bands, principally the alpha, beta, gamma, and high gamma bands. Grey bars at the foot of each plot indicate Chance-level participant differentiation accuracy computed using ’mock brain-fingerprints’ derived from environmental and instrument noise recorded during MEG sessions. Error bars represent 95% confidence intervals.

**Figure S2.**
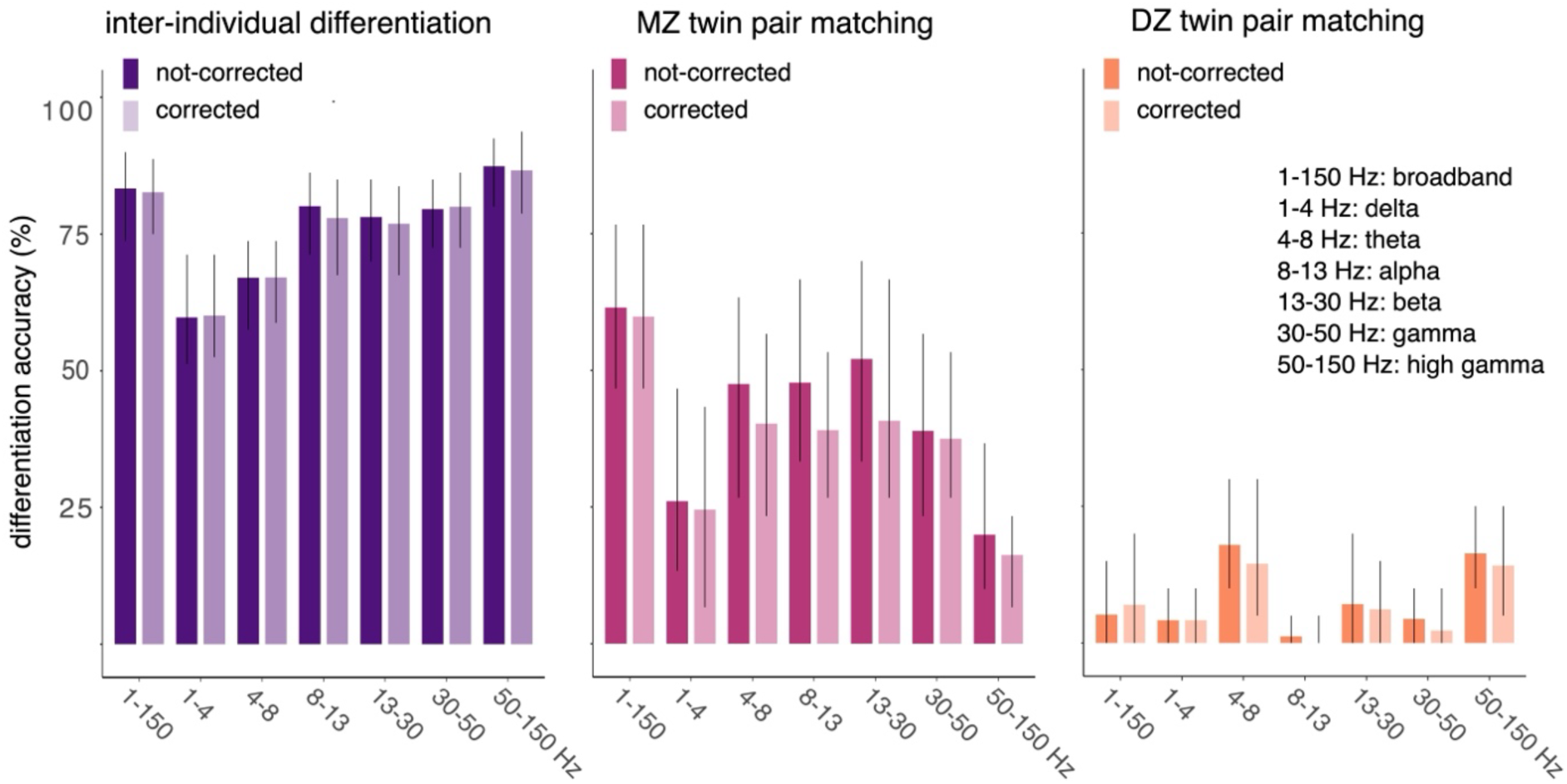
Physiological artifacts do not impact differentiation & matching accuracy. Comparison of the differentiation and twin-matching accuracy scores before and after regressing out the influence of artifacts on neurophysiological profiles, for singletons (left panel), MZ twin pairs (middle panel) and DZ twin pairs (right panel).

**Figure S3:**
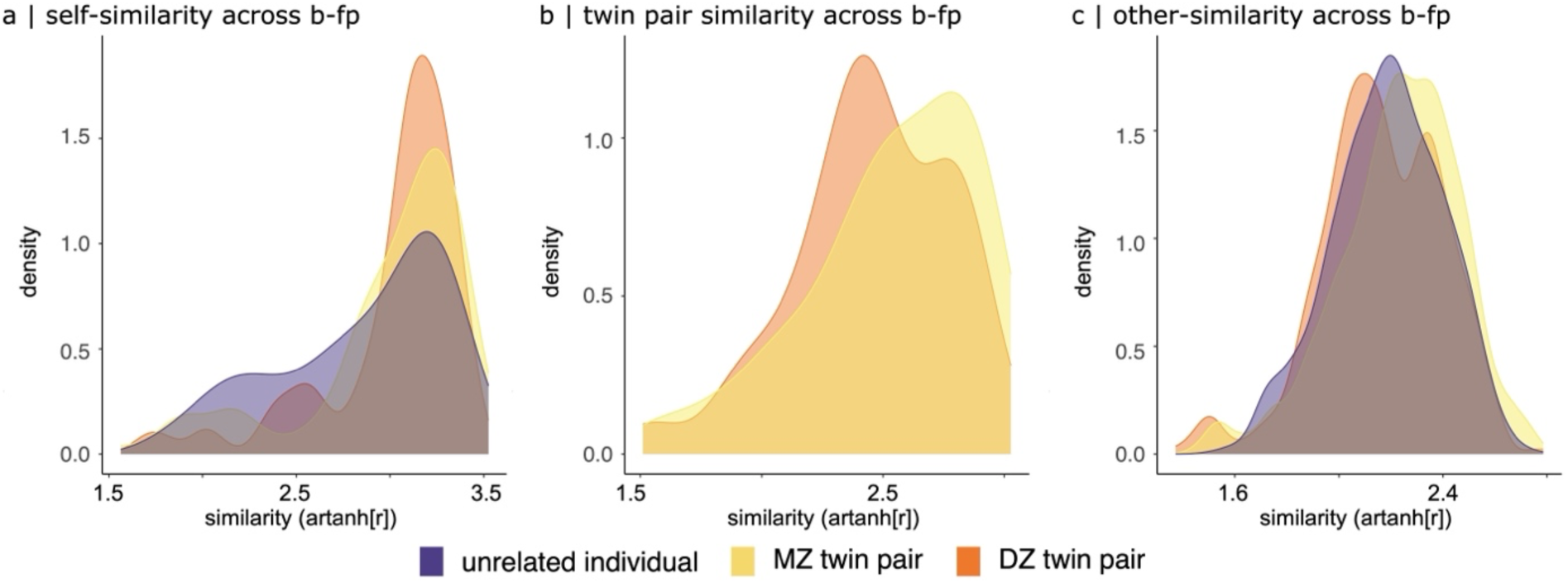
Self-, twin-pair, and other-similarity between neurophysiological profiles. a) Self-similarity did not significantly differ between monozygotic (MZ) and dizygotic (DZ) twin pairs. Unrelated individuals exhibited a similar mean self-similarity but greater inter-individual variability (higher standard deviation). b) Neurophysiological profiles (1–150 Hz) within MZ twin pairs were more similar than those of DZ twin pairs, and twin-pair similarity was greater than other-similarity. c) Similarity between neurophysiological profiles of unrelated individuals (other-similarity) did not significantly differ across the three groups (MZ & DZ twins, and non-twin individuals). Legend: b-fp, brain-fingerprint.

### The Matching Between the Neurophysiological Profiles of Monozygotic Twins Is Not Driven by Anatomy

We tested whether the matching between the neurophysiological profiles of MZ twins may have been driven by similarities in their brain anatomy. The anatomical features extracted included: the Number of vertices, surface area, gray matter volume, mean cortical thickness, s.d. of cortical thickness, mean curvature, Gaussian curvature, folding index, and the curvature index (*9*). The brain structural features of MZ siblings extracted from *Freesurfer* (*9*) were indeed more similar than those of DZ twins or unrelated participants (see Methods): they showed a high correlation between MZ siblings (r= 0.99), and lower for DZ twin pairs (r= 0.93). The most heritable brain anatomical features the most heritable were related to the curvature (h= 1.75) and thickness (h= 1.75) of the cortex.

To assess the extent to which these brain structural features contributed to the heritability of neurophysiological profiles, we estimated the linear correlation between the matching of neurophysiological profiles of twin siblings and the similarity of their respective brain anatomical features (see Methods). We matched twin pairs using all 9 features for every parcel of the Desikan-Killiany atlas. These relationships were not statistically significant between MZ siblings (r= 0.32, p= 0.06) nor between DZ siblings (r= -0.22, p= 0.32; see Table S2). The outcome was similar when we replicated this analysis for neurophysiological profiles derived from alpha-band (MZ: r= 0.31, p= 0.08; DZ: r= -0.17, p= 0.45) and beta-band (MZ: r= 0.32, p= 0.07; DZ: r= -0.15, p= 0.51) brain activity. Bayes factor analyses corroborated that there was little evidence for a relationship between similarities of structural and neurophysiological traits (Supplemental Table S2). To conclude, while brain curvature and cortical thickness are heritable brain phenotypes, they did not contribute significantly to individual differentiation based on their neurophysiological profiles.

**Table S2:** Pearson’s Correlations Between the Similarity of Brain Anatomy and Neurophysiological Profiles.

| | Pearson's $r$ | | $BF_{10}$ | |
| --- | --- | --- | --- | --- |
|  | MZ | DZ | MZ | DZ |
| Anatomy vs. broadband neurophysiology | 0.32 | -0.22 | 1.65 | 0.68 |
| Anatomy vs. alpha-band neurophysiology | 0.31 | -0.17 | 1.46 | 0.57 |
| Anatomy vs. Beta-band neurophysiology | 0.31 | -0.15 | 1.60 | 0.54 |

We fit linear models relating the similarity of brain anatomical similarity and neurophysiological profiles derived from broadband (1-150 Hz), alpha band, and beta band activity between MZ and DZ twin siblings. There was little Bayes factor evidence (BF_10_) of such a relationship.

### Salient Features for Participant Differentiation are Heritable

**Figure S4.**
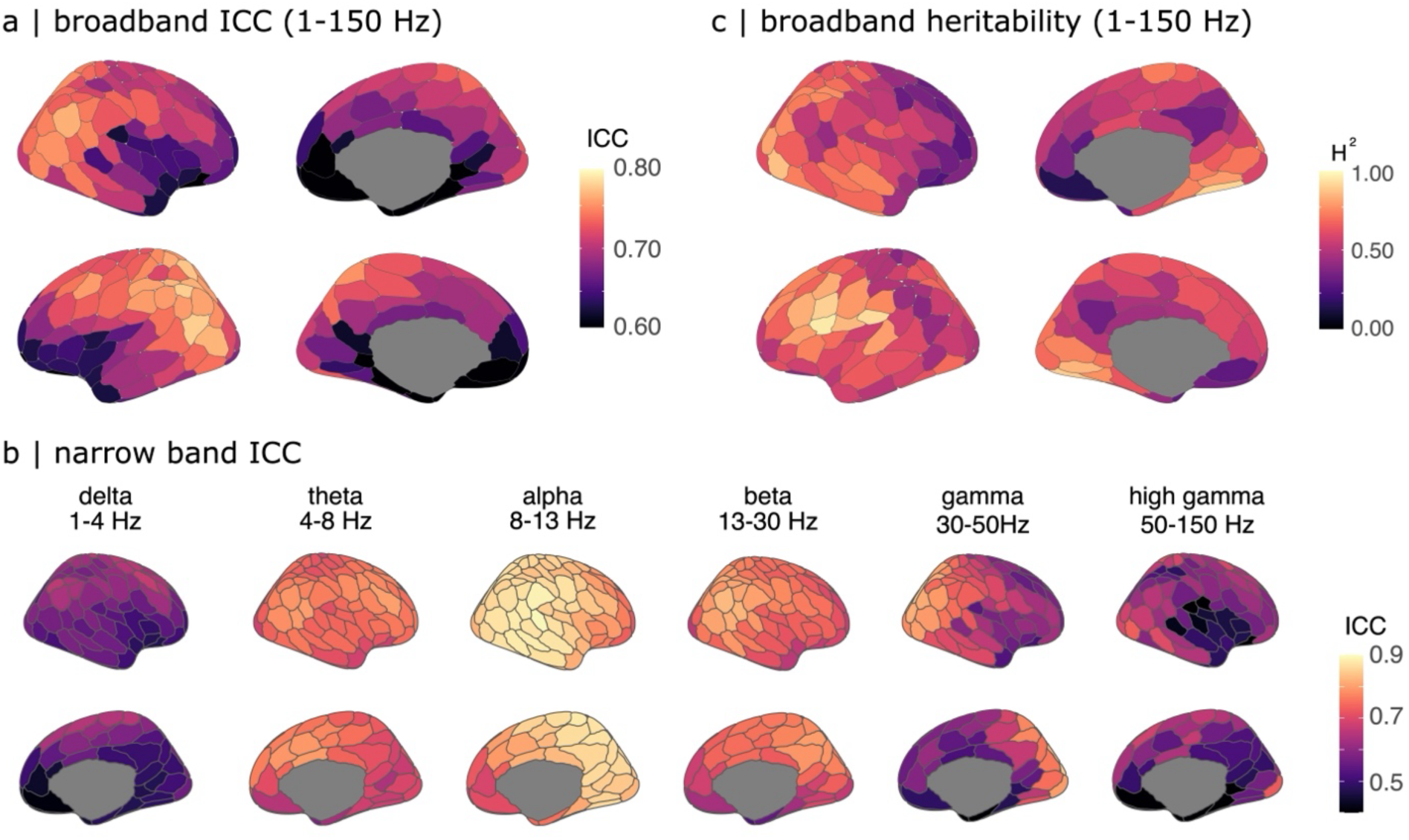
Salient Regions of Neurophysiological Profiles and Their Heritable Brain Phenotypes. (a & b) Topographic maps highlight the cortical regions with the most salient neurophysiological activity across all frequency bands (panel a) and per frequency band (panel b), as measured using intra-class correlation (ICC) statistics (see Main Text and Methods). These regions contribute most strongly to participant differentiation. Note that the broadband (1-150 Hz) ICC topographic maps in panel (a) were obtained by averaging the ICC values across the six narrow frequency bands (delta, theta, alpha, beta, gamma, high gamma), ensuring equal weighting regardless of bandwidth differences. (c) Topographic maps depicting the heritability of broadband neurophysiological activity (1-150 Hz). For frequency-specific maps of heritable neurophysiological traits, refer to Figure 1C in the main text.

**Figure S5.**
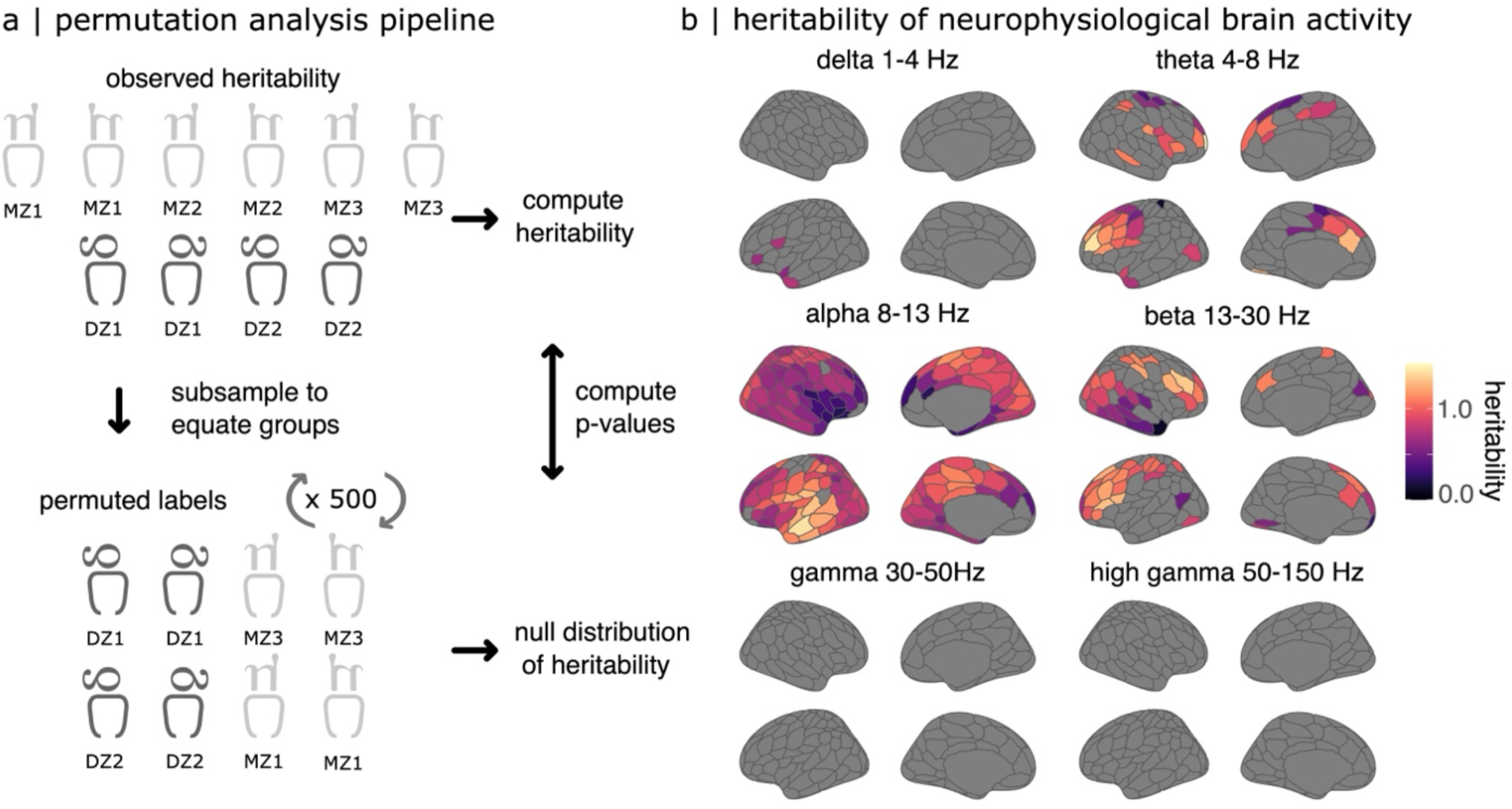
Significantly Heritable Neurophysiological Traits. a) Analysis pipeline for assessing the statistical significance of heritability estimates. Heritability (H^2^) was computed as the difference in neurophysiological profile concordance between monozygotic (MZ) and dizygotic (DZ) twin pairs. To generate a spatially realistic null distribution, MZ and DZ twin labels were randomly reassigned across 1000 iterations, after first subsampling MZ pairs to match the number of DZ pairs in each permutation. Spatial autocorrelation of the data was preserved throughout. b) Results of the permutation analysis across frequency bands and cortical regions. Cortical maps show brain regions where heritability estimates significantly exceeded chance levels (p_FDR_ < 0.05). Significant heritability was observed in frontal regions for theta-band activity, across widespread cortical areas for alpha-band activity, and in frontal and parietal regions for beta-band activity.

**Figure S6:**
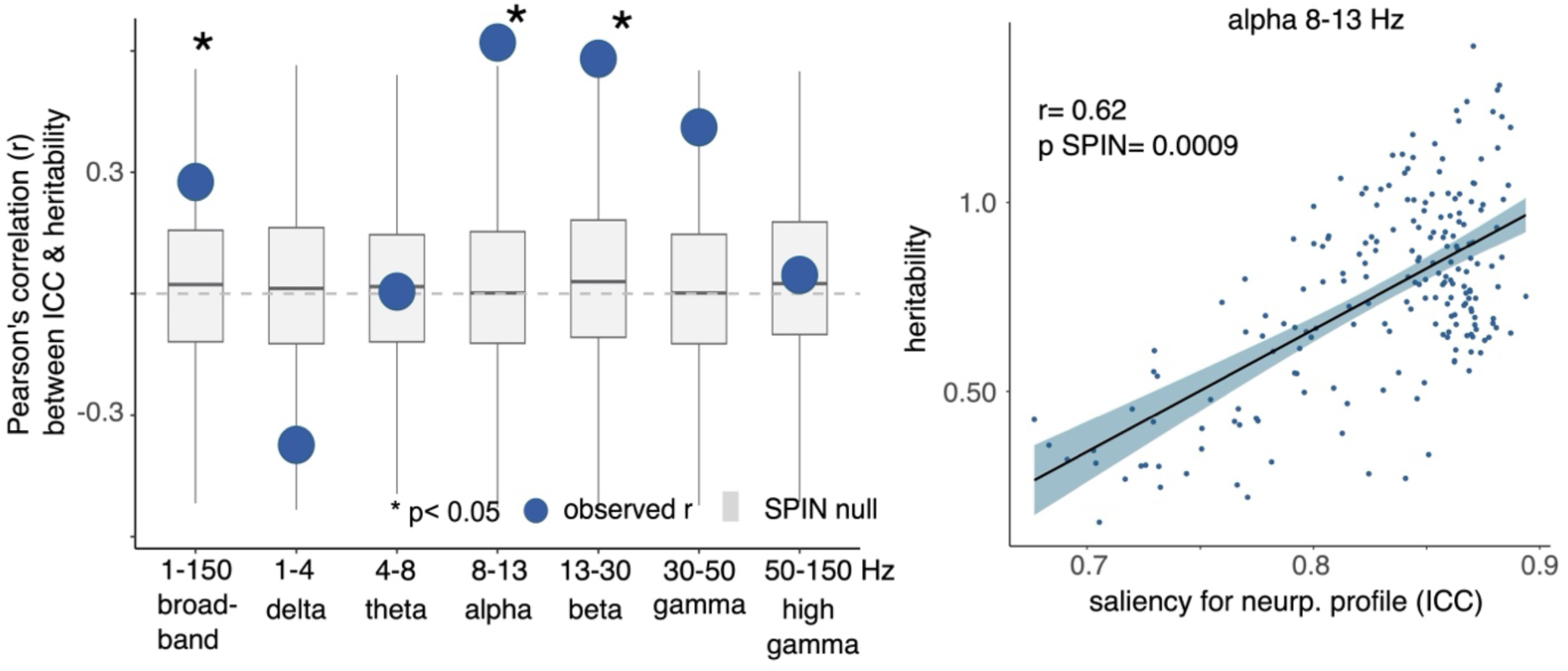
Differentiable Neurophysiological Traits are Heritable. <u>Left panel:</u> Pearson’s correlation between the neurophysiological traits that are the most salient for individual differentiation and their heritability, for each tested frequency band of electrophysiology. The blue dots indicate the correlation statistics and the boxplots depict the null distributions obtained by spin-test permutations. <u>Right panel:</u> Scatter plot of best linear model relating the heritability and saliency of alpha-band neurophysiological traits. Each dot represents a region of the Schaefer-200 atlas. The saliency of the alpha-band neurophysiological traits for individual differentiation is linearly related to their heritability. This further demonstrates the genetic influence on neurophysiological traits.

#### Psychological Processes and Differentiation

We assessed the similarity in gene-expression brain score between the outcomes of the gene-differentiation and gene-psychological processes PLS analyses. We found strong linear relationships between the identified gene scores and the PLS loadings (The Gene-Differentiation Gradient Correlates with Psychological Processes).

We further tested whether the outcome of the PLS analysis for psychological processes (see Main Text) covaried with individual differentiation. We anticipated such a relationship as both psychological processes and individual differentiation covary with similar gene expression signatures. The PLS of psychological-process and individual differentiation featured a single significant latent variable (p= 0.002) that explained 87.0% of the covariance between these variables (85.2% covariance explained, p_SPIN_= 0.002, 95% CI = [54.24., 87.37]). The ICC loadings and psychological process term loadings were linearly related to the loadings obtained from the previously reported PLS analysis (ICC loadings similarity, r=0.78; term loading similarity, r= 0.92).

## Supplementary Data

Supplemental Data 1: Negative gene set used for the gene ontology analysis. Table of the gene name, EntrezID, and PLS loading of the negative set of genes used in the GO analysis.

Supplemental Data 2: Positive gene set used for the gene ontology analysis. Table of the gene name, EntrezID, and PLS loading of the positive set of genes used in the GO analysis.

Supplemental Data 3: Results of the gene ontology analysis for the negative gene set. Table of the GO results for the negative gene set. Rows correspond to biological processes from the GO analysis, with their corresponding fold enrichment, p-value, and the corresponding genes that make up that GO category.

Supplemental Data 4: Results of the gene ontology analysis for the positive gene set. Table of the GO results for the positive gene set. Rows correspond to biological processes from the GO analysis, with their corresponding fold enrichment, p-value, and the corresponding genes that make up that GO category.

